# Determination of m^6^A frequency utilizing 4SedTTP-RT Ligation Assisted PCR (SLAP) in viral and cellular long non-coding RNAs

**DOI:** 10.1101/2021.09.16.460679

**Authors:** Sarah E Martin, Huachen Gan, Joanna Sztuba-Solinska

## Abstract

N^6^-methyladenosine is one of the most abundant epitranscriptomic signatures that can affect every aspect of RNA biology, from structure and stability to intra- and intermolecular interactions. The accurate quantitative assessment of RNA stoichiometry at single-nucleotide resolution is a prerequisite to evaluate the biological significance of m^6^A in the context of specific RNA. We have developed a new method, termed 4-<u>S</u>elenothymidine 5’-triphosphate reverse transcription and <u>L</u>igation <u>A</u>ssisted <u>P</u>CR analysis (SLAP), for quantitative and unbiased assessment of the m^6^A fraction on target RNA. The inclusion of thymidine triphosphate derivative during reverse transcription discourages base pair formation with m^6^A resulting in the reaction’s cessation, while maintaining normal A-T base pairing. The site-specific ligation of the resulting cDNAs with adapters, followed by amplification, generates two distinct products that reflect the modified and unmodified fraction of the analyzed RNA. These PCR products are subsequently separated by gel electrophoresis and quantified using densitometric analysis. We applied the SLAP to verify the position and assess the frequency of m^6^A sites present on two exemplary long non-coding RNAs. We assessed the SLAP specificity, accuracy, and sensitivity, proving the applicability of this method for the m^6^A analysis on less abundant transcripts. Overall, this method constitutes an extension of the bird’s-eye view of RNA m^6^A landscape provided by epitranscriptome-wide analyses by delivering quantitative assessment of modification frequency and can therefore aid the understanding of the consequences of m^6^A on biological processes.

**Graphical Abstract:** 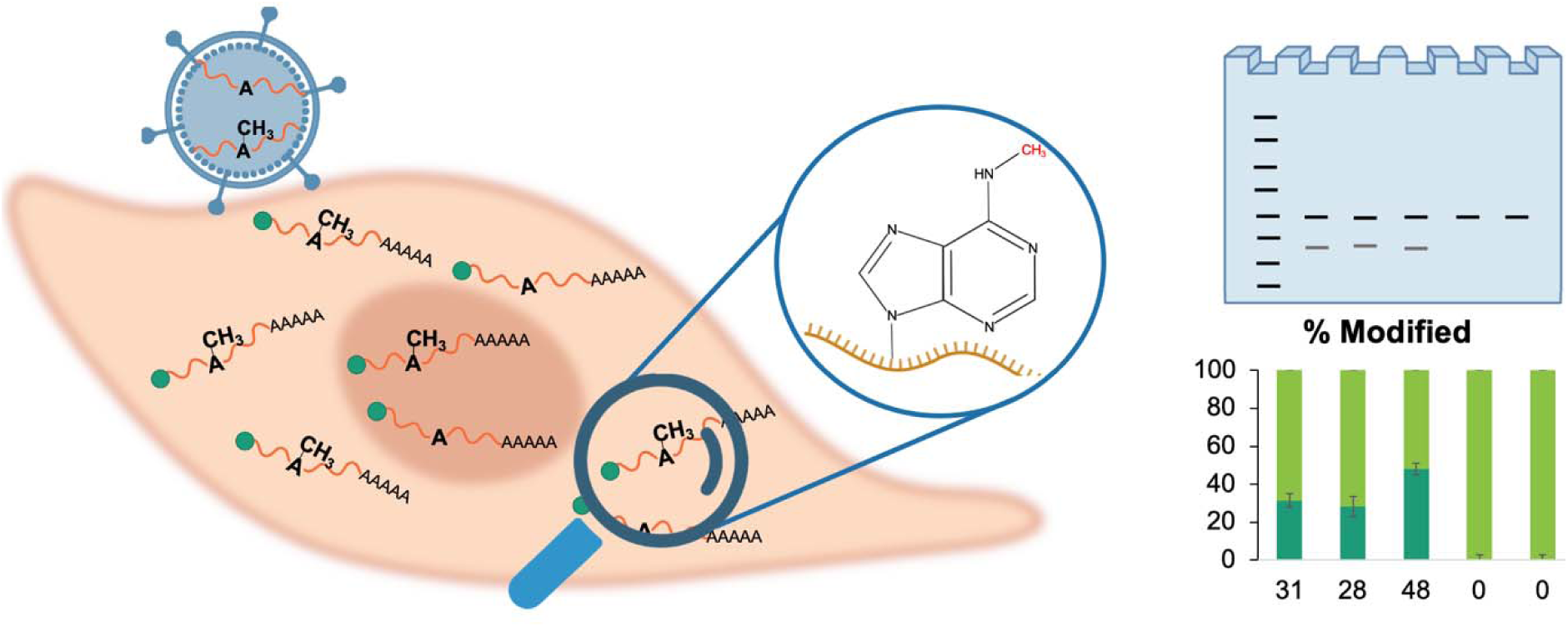

## Background

N^6^-methyladenosine (m^6^A) is an epitranscriptomic modification that involves the methylation of the sixth position of the base moiety of adenine. It is one of the most abundant chemical signatures found on coding and non-coding (nc) RNAs expressed by all living taxa (Jiang et al. 2021; Coker et al. 2019; Liu et al. 2014; Bokar et al. 1997; Wang et al. 2016), and non-living forms, i.e., viruses (Wu et al. 2019; Tan and Gao 2018; McIntyre et al. 2018; Baquero-Perez et al. 2021). For the past few decades, the biological significance of m^6^A remained elusive. Recent developments in the next-generation sequencing methods brought the long-awaited breakthrough in the field and demonstrated the overwhelming significance of m^6^A for RNA biology. It has been shown that m^6^A functions at almost all stages of messenger (m) RNA lifetime, including splicing, translation, stability, export, and subcellular localization (Zhou et al. 2019b; Bartosovic et al. 2017; Zhou et al. 2019a; Meyer et al. 2015; Wang et al. 2015, 2014; Kane and Beemon 1985). For example, the installation of m^6^A within the splice site of precursor mRNA coding for S-adenosylmethionine synthetase inhibits its proper splicing and translation (Mendel et al. 2021). Also, m^6^A found within the coding region of mRNAs positively regulates translation by resolving RNA secondary structures (Mao et al. 2019). In ncRNAs, the breadth of m^6^A impact is attributed to local and global structural changes that can influence the accessibility of RNA motifs for effectors binding. For example, the modification of metastasis-associated lung adenocarcinoma transcript 1 (MALAT1) regulates the binding of heterogeneous nuclear ribonucleoprotein C (HNRNPC) to U-rich hairpin. Likewise, the modification of A-repeat domain in X-inactive specific transcript (XIST) facilitates the specific folding of the transcript, which guides proteins involved in transcriptional silencing to their RNA binding motifs (Lu et al. 2020).

The m^6^A is a highly dynamic signature regulated by the action of methyltransferases and demethylases. Methyltransferases, referred to as “writers”, install the m^6^A modification. Demethylases act as “erasers,” remove the chemical tag (Yang et al. 2018; Schwartz et al. 2014). The writers are thought to recognize the consensus sequence RRACH (where R is a purine, H is adenine, cytosine, or uracil). Not every RRACH motif, however, undergoes methylation (Bokar 2005). This fact suggests that RNA structural determinants can influence the deposition of this chemical tag (Mateusz Mendel et al. 2018). Most m^6^A are installed by METTL3/14 complex (Śledź and Jinek 2016), the binding and catalytic activity of which seems to be independent of substrate’s structure, while the activity of another writer, METTL16, is structure mediated (Mateusz Mendel et al. 2018). The direct outcome of writer and eraser activity influences the position, frequency, and the overall abundance of m^6^A, thus determining the breadth of its impact on RNA biology. For example, suppose methylation at a given site serves as a gene regulatory mechanism. In this instance, the protein occupancy of controlled modification site is expected to vary over the lifetime of RNA to alter its function.

Despite the rapidly growing recognition of m^6^A significance, the prevalence and functional consequences of m^6^A remain elusive as the currently available detection methods are primarily qualitative. The widely available antibody-based next-generation sequencing mapping methods, including m^6^A-seq (Dominissini et al. 2012), methylated RNA immunoprecipitation (m^6^A-MeRIP) (Meyer et al. 2012), and m^6^A individual-nucleotide-resolution crosslinking and immunoprecipitation (miCLIP) (Linder et al. 2015), are burden with various shortcomings, e.g., the high signal-to-noise ratio, large number of false positives due to cross-reactivity of antibodies with related modifications (Linder et al. 2015; Dominissini et al. 2012). Furthermore, since these methods involve the multi-step protocols for library preparation, they often result in poor data reproducibility, hindering the between-sample analyses. Site-specific cleavage and radioactive-labeling followed by ligation-assisted extraction and thin-layer chromatography (SCARLET) uses complementary oligonucleotides targeted to known modification sites to investigate the occupancy of the modification but is an exceptionally laborious method that relies on radioactivity (Liu et al. 2013a). The <u>L</u>iquid <u>C</u>hromatography with tandem <u>m</u>ass <u>s</u>pectrometry (LC-MS/MS) is currently the only technology that can directly and comprehensively estimate the frequency of epitranscriptomic modifications. However, the information about the sequence context and co-occurrence of other modifications is lost during RNA hydrolysis, and the method demands large quantities of highly purified RNA (Wein et al. 2020).

In this manuscript, we provide a detailed outline of a novel antibody-independent termination-based method for the target-specific quantification of m^6^A frequency (Figure 1). Our method, termed <u>S</u>elenium-modified deoxythymidine triphosphate reverse transcription and <u>L</u>igation <u>A</u>ssisted <u>P</u>CR (SLAP), relies on the use of 4-selenothymidine-5’-triphosphate (4SedTTP) during reverse transcription (RT). 4SedTTP carries a selenium atom at the 4-position of deoxythymidine triphosphate instead of oxygen. That replacement affords exclusive hybridization properties when incorporated into TTP nucleobase due to its unique steric and electronic efforts (Hong et al. 2018). During RT, the 4SedTTP supports A-T pairing and at the same time discourages m^6^A/T pairing, triggering RT termination at +1 position from the modification (Hong et al. 2018). Truncated and full-length products corresponding to modified and unmodified adenines, respectively, are subsequently ligated to an adapter carrying a primer binding site to allow for the simultaneous and uniform amplification of both types of cDNA products in a single PCR reaction. This approach yields quantitative and unbiased information for each m^6^A site, and because the SLAP method relies on the use of target- and site-specific RT primers, the modification frequency can be determined regardless of its location, i.e., inside, or outside of the consensus sequence.

**Figure 1.**
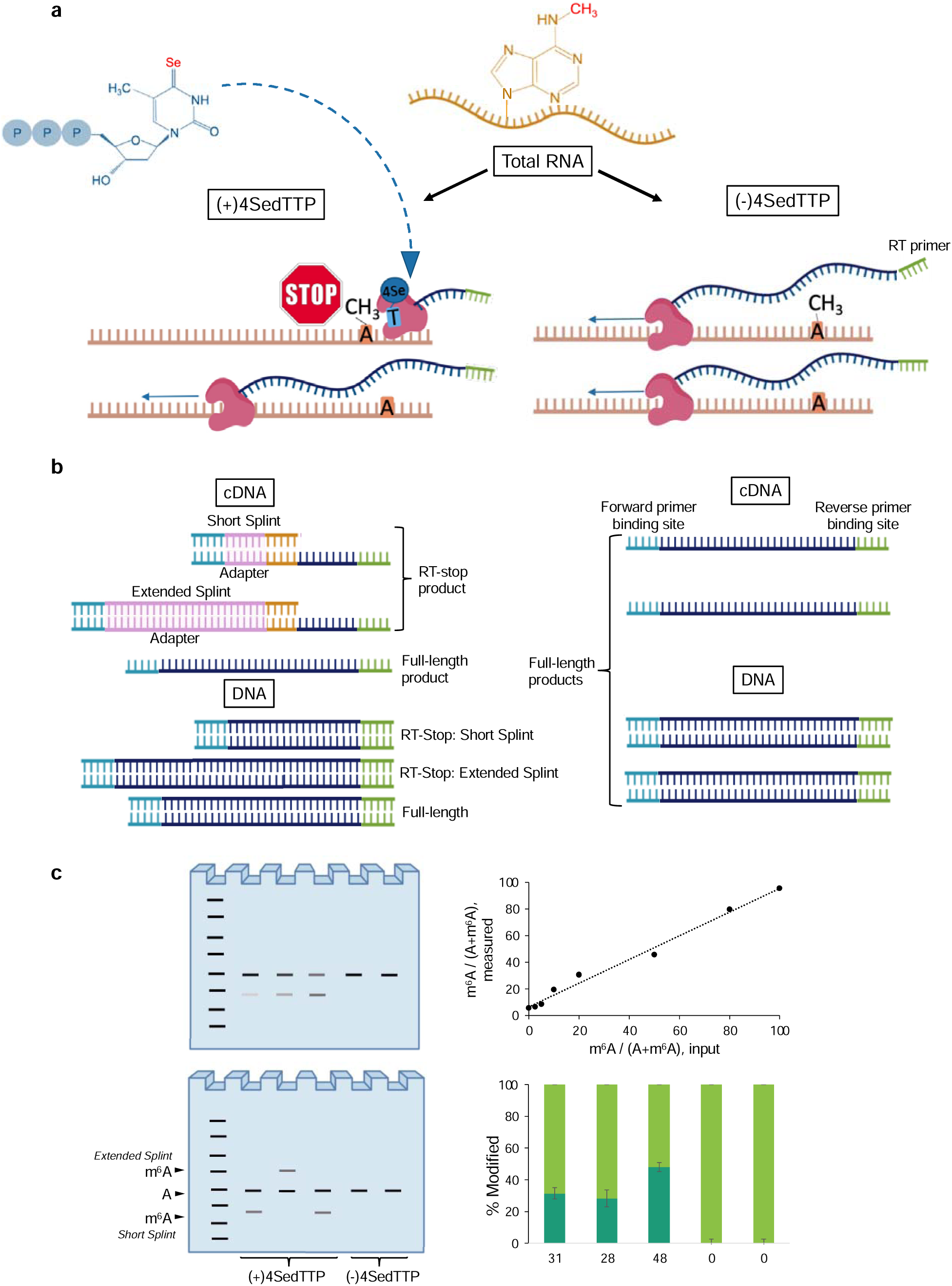
Schematic overview of the SLAP method. a, Reverse Transcription. 4SedTTP is used during reverse transcription reaction (RT) to prompt RT-stop formation at +1 position from m^6^A. **b, Splint bridge and adapter ligation**. A splint oligonucleotide is annealed to the truncated cDNA product that results from reverse transcription of the modified RNA. The splint oligonucleotide has a 5’ complementarity to the adapter oligonucleotide that is ligated to RT-stop product to introduce the reverse primer binding site. For m^6^A close to the 5’ or 3’ end of the transcript, extended splint should be utilized to allow for size-differentiation between modified and unmodified products. **C, Densitometric analysis of experimental results**. Subsequent PCR reaction performed using a common set of forward and reverse primers, simultaneously amplifies both full-length and RT-stop products. The PCR products are visualized on native polyacrylamide gel (PAGE) and directed to densitometric analysis. The densitometric measurements are compared to the established calibration curve to assess the m^6^A stoichiometry.

We applied SLAP to analyze the frequency of m^6^A at multiple sites found on two long non-coding (lnc) RNAs, i.e., polyadenylated nuclear (PAN) RNA encoded by Kaposi’s sarcoma-associated herpesvirus (KSHV) (Martin et al. 2021), and cellular lncRNA MALAT1 (Yang et al. 2013). We assessed the sensitivity of this method by titrating in vitro synthesized PAN transcript to the total cellular RNA and revealed that the method provides an accurate estimation of modification frequency at an attomolar concentration of the target. This highlights the potential of the SLAP method for analyzing m^6^A stoichiometry on low abundance RNAs. Paired with any of the available next-generation sequencing methods, our protocol not only provides further verification of m^6^A localization at the single-nucleotide resolution but, most importantly, the quantitative estimate of modification abundance.

## RESULTS

### Target m^6^A modified RNAs

PAN RNA is a lncRNA expressed by Kaposi’s sarcoma-associated herpesvirus (KSHV) at low levels during latency (10^3^ copies/cell according to our estimates, data not published), but it can reach up to 10^5^ copies/cell 24 hours post lytic induction in BCBL-1 cells. PAN RNA was proposed to associate with chromatin modulating complexes, histones, and has been implicated in the altering of viral and cellular gene expression (Rossetto and Pari 2014). Other studies suggest that PAN RNA facilitates late viral mRNA export from the nucleus to the cytoplasm (Withers et al. 2018). We recently applied 4-selenothymidine-5′-triphosphate reverse transcription (4SedTTP RT) with next generation sequencing method to analyze the PAN RNA m^6^A landscape during KSHV replication (Martin et al. 2021). Our findings showed that PAN RNA can carry up to five m^6^A residues during the late lytic stages of KSHV replication. The functionality of these residues is yet to be determined.

MALAT1 is a nuclear lncRNA which exhibits copy number changes (on average 2,500 copies/cell (Tripathi et al. 2010), translocations, or mutations in several cancer types (Arun et al. 2020). MALAT1 was shown to be m^6^A modified by the application of m^6^A-specific methylated RNA immunoprecipitation with next-generation sequencing (m^6^A/MeRIP-Seq) (KD et al. 2012). Since the m^6^A/MeRIP-seq method combines m^6^A antibody immune-precipitation and deep sequencing to locate m^6^A in ∼200 nt RNA segments, it cannot identify which adenosine residue under the deep sequencing peaks is modified, nor can it determine the modification fraction for any modification site. To address this challenge, Liu et al. applied the SCARLET method to directly measure the location and m^6^A fraction on MALAT1 in three different cell lines (Liu et al. 2013a). They found that m^6^A is present at four out of seven previously identified sites, and that, the modification fraction varied by up to threefold between cell lines. The m^6^A sites on MALAT1 were shown to regulate protein binding through invoking the RNA structural changes. In particular, m^6^A modification at nt 2577 was shown to destabilize the modified hairpin and release a poly-U tract for an increased binding of m^6^A reader protein, Heterogenous Nuclear Ribonucleoprotein C (HNRNPC) (Yang et al. 2013; He et al. 2020).

### Establishing the SLAP calibration curves for target RNAs

To estimate the m^6^A stoichiometry on target RNA, it is necessary to establish the proper calibration curve. We combined 100 nt in vitro synthesized m^6^A modified (nucleotide (nt) position 82 on PAN RNA and nt position 65 on MALAT1) and unmodified RNA standards at the following ratios (0:1, 40:1, 20:1, 9:1, 4:1, 1:1, 1:4, and 1:0), that reflected the modification percentage (0, 2.5, 5, 10, 20, 50, 80, and 100%, respectively) for the total of 30 femtomole, and spiked them with the 1 µg total RNA extracts. This approach is meant to mimic the experimental conditions one would work with during the SLAP analysis performed on RNA expressed in living cells. The combined RNA standards were directed to reverse transcription reactions (RT) using the avian myoblastoma virus (AMV) reverse transcriptase. Other types of reverse transcriptase, including SuperScript II and III, perform equally well in the 4SedTTP RT reactions, and as such they can replace AMV (Hong et al. 2018; Martin et al. 2021). Each ratio of combined RNA standards was directed to two RT reactions, i.e., positive reaction performed in the presence of SedTTP (+SedTTP RT) and negative control reaction, in which SedTTP was replaced with TTP (-SedTTP RT). The RT-stop cDNA products resulting from reverse transcription of modified RNAs were site-specifically ligated with splint-adapter oligonucleotide duplex to yield products that can be simultaneously amplified with full-length products that correspond to unmodified RNA. Following amplification, the products were resolved on native polyacrylamide gel and quantified.

From the calibration curve established for PAN RNA, we estimated that the SLAP allows for quantification of m^6^A frequency at as low as 2.5% level (4.5 × 10^8^ copies modified). The positive reactions that included unmodified transcript (+0%, Figure 2a) yielded a weak m^6^A-specific product, which was considered as the background, and as such a 2-fold threshold above that background was applied to assess the modification stoichiometry. Also, a minor background product was notable for negative reactions, suggesting the occurrence of non-specific-to-m^6^A truncations in some RT reactions. Interestingly, the sample including 100% modified transcript resulted in an estimation of m^6^A frequency at 92% level, which suggests a minor underestimation of the actual modification fraction. Overall, for these combined standards we achieved linear regression of R^2^ = 0.988 (Figure 2b). Using the same approach, we also established a calibration curve for MALAT1 (Figure 2d), which allowed for the quantification of m^6^A frequency at equally low levels (2.5%) compared to PAN RNA, and we obtained the linear regression of R^2^= 0.985 (Figure 2e). The establishment of an optimized standard curve with an R^2^ value >0.980 is critical for the precise estimation of m^6^A modification frequency.

**Figure 2.**
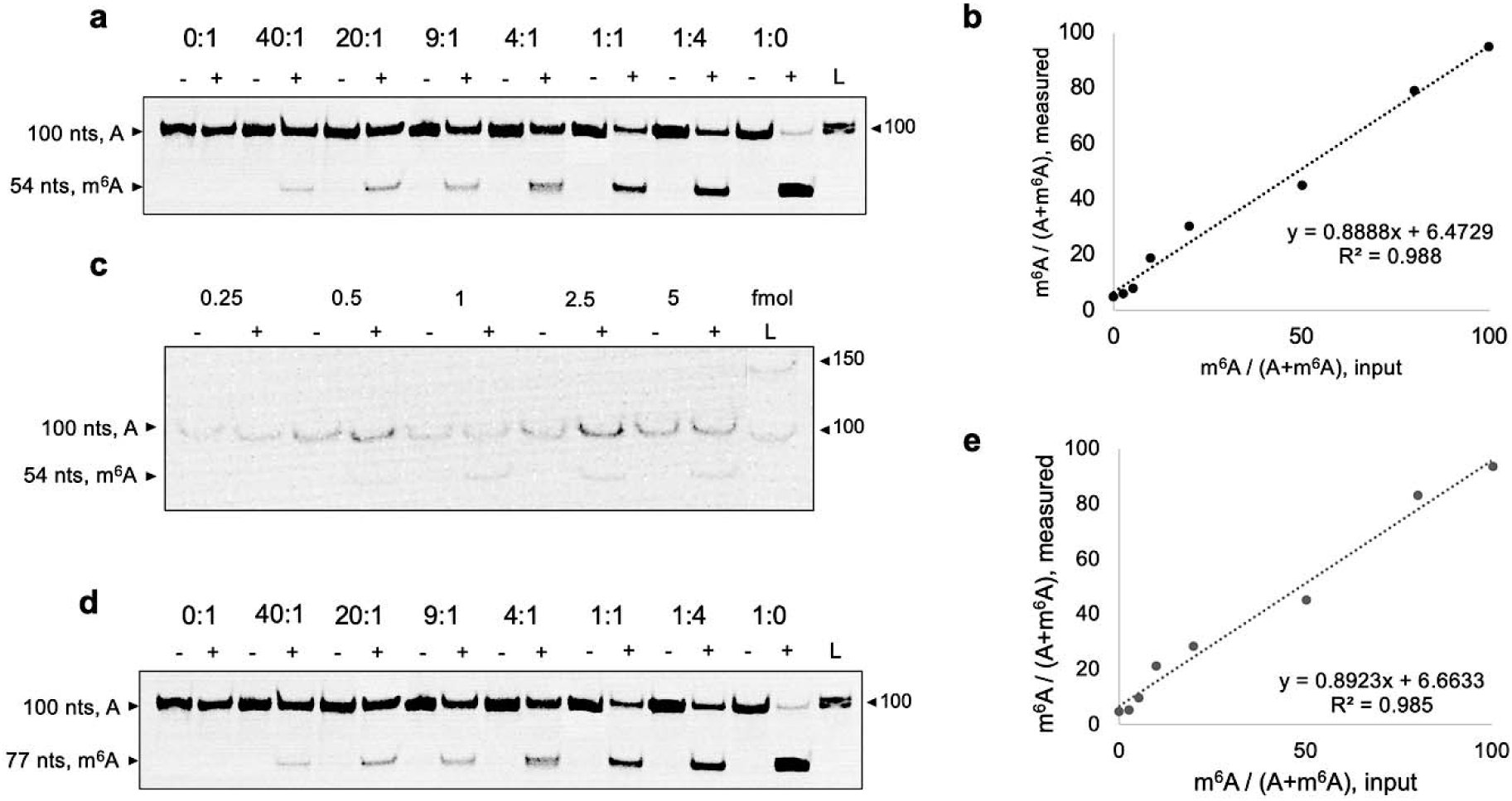
The calibration curves for SLAP analysis of PAN and MALAT1 lncRNAs. **a**, To estimate the m^6^A stoichiometry on target RNA, it is necessary to establish the proper calibration curve. We combined 100 nt in vitro synthesized m^6^A modified (nucleotide (nt) position 82 on PAN RNA and nt position 65 on MALAT1) and unmodified RNA standards at the following ratios (0:1, 40:1, 20:1, 9:1, 4:1, 1:1, 1:4, and 1:0), that reflected the modification percentage (0, 2.5, 5, 10, 20, 50, 80, and 100%, respectively) for the total of 30 femtomole, and spiked them with the 1 µg total RNA extracts. Native polyacrylamide gel (PAGE) shows two products corresponding to the m^6^A modified (54 nts) and unmodified (100 nts) PAN RNA standards that were combined at indicated ratios and directed to SLAP analysis. Positive reactions included SedTTP in RT reactions, while negative reactions included dTTP instead. The intensity of these products was quantified to generate the calibration curve shown in panel **b**, which showed linear regression of R^2^ = 0.988. **c**, Native PAGE showing two products corresponding to the modified (54 nts) and unmodified (100 nts) standard RNAs that were combined at equal ratio at the indicated total concentrations (0.25 – 5 femtomole) and directed to the SLAP analysis to assess the method’s sensitivity. **d**, Native PAGE gel showing two products derived from the m^6^A modified (77 nts) and unmodified (100 nts) MALAT1 RNA standards that were combined at specific ratios and directed to the SLAP analysis. **e**, The intensity of the products from panel d was quantified to generate the calibration curve, which showed linear regression of R^2^ = 0.89.

To test the sensitivity of the SLAP method, we performed a serial dilution of combined at 1:1 ratio m^6^A modified and unmodified RNA standards, that were subsequently spiked with 1 µg of total RNA and directed to the analysis. We tested the total concentrations of 0.25, 0.5, 1, 2.5, and 5 femtomole RNA to determine the minimum concentration that allows for the determination of m^6^A frequency. We were able to quantify the stoichiometry of m^6^A on the target RNA that was present at as low as 500 attomolar (aM) concentration (5.5 × 10^8^ copies, Figure 2c). Thus, the SLAP can likely be applied to analyze the frequency of m^6^A on low abundance target RNAs.

### Stoichiometric analysis of m^6^A on PAN RNA

After establishing the proper calibration curve, we proceeded to the quantification of m^6^A frequency on PAN RNA at three selected nt positions, i.e., 18, 203 and 1041, which, according to our previous next generation epitranscriptomic analysis, are modified during uninduced (0 hours post-induction, h pi) and late lytic (48 h pi) stages of KSHV replication (Martin et al. 2021). We included the analysis of two unmodified adenines on PAN RNA, at nt positions 366 and 410, as negative controls (Figure 3a). Here, the analysis did not yield m^6^A-specific products, verifying the SLAP specificity.

**Figure 3.**
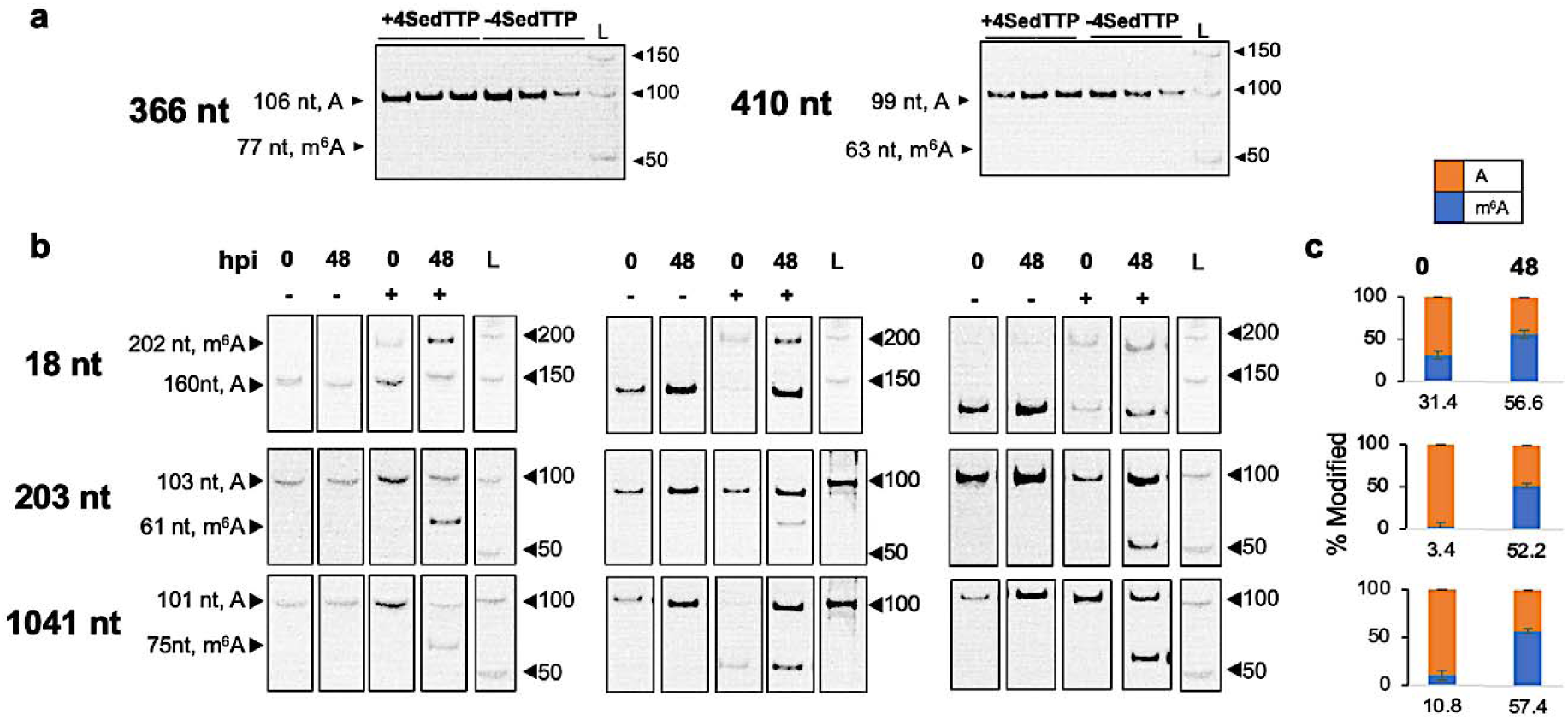
The SLAP analysis of m^6^A stoichoimetry on PAN RNA. **a**, Native PAGE showing one product corresponding to the unmodified adenines at nt position 366 and 410 that served as negative controls. The position of expected products specific for modified and unmodified sites are indicated on all electropherograms. L stands for 50 bp DNA ladder. **b**, Native PAGE for representative three biological replicates of the SLAP analysis performed on PAN RNA for adenines at nt positions 18, 203, and 1041. **c**, The column graphs represent the average m^6^A modification frequency at nt positions 18, 203, and 1041. Standard deviations for frequency measurements are indicated.

Due to the proximity of m^6^A at nt position 18 to the 5’ terminus, analysis of that site required the use of extended splint-adapter oligonucleotide duplex containing the forward primer binding site (extension follows the CCATTG insert and proceed the 3’ end sequence that is reverse complement to the forward primer, see Materials and Methods section for details), that would allow size-specific differentiation of modified and unmodified products. As a result, the m^6^A-specific product corresponding to that site is 42 nts longer (the total length of 202 nts) than the product corresponding to unmodified residue (160 nt) (Figure 3b). For nt positions 203 and 1041, the size of the products corresponding to modified and unmodified RNA fractions was 103 and 75, respectively (Figure 3b).

The m^6^A at position 18 was estimated at 31.4±3.5% during uninduced stage of KSHV replication, followed by an increase to 56.6±2.7% during late lytic phase of infection (Figure 3c). The modification level at nt 203 was estimated at 3.4±4.8% during uninduced stage, followed by an increase to 52.2±5.2% during the late lytic stage (Figure 3c). Nucleotide 1041 was modified at 10.8±5.2% during uninduced stage and showed the highest modification frequency during the late lytic stage, reaching a frequency of 57.4±5.3% (Figure 3c).

### Stoichiometric analysis of m^6^A on MALAT1

We used the SLAP analysis to estimate the modification frequency of two previously identified m^6^A at nt 2515 and 2698 on MALAT1 expressed during uninduced (0 h pi) and late lytic (48 h pi) phases of KSHV replication. We included the analysis of unmodified adenine at nt position 2674 as a control (Figure 4a). Previous application of the SCARLET analysis indicated that MALAT1 is modified at nt 2515 at 41% frequency in HEK293T cells (Liu et al. 2013b). In our studies, we found that the modification frequency of this position is dynamic and varies depending upon KSHV replication stage. During the uninduced stage, nt 2515 was modified at 9.28±8.28% frequency, however, the progression of viral lytic replication led to the increased m^6^A frequency to 20.39 ±6.78% in BCBL-1 cells (Figure 4c). The SCARLET analysis of another site on MALAT1 at nt 2698 indicated the modification frequency at 3.77±0.23%. In the SLAP analysis, this position was found to be modified at 3.77±0.23% during the uninduced stage of KSHV replication, while during lytic replication the frequency increased to 7.76±0.54% in BCBL-1 cells (Figure 4c). The observed varying levels of m^6^A on both analyzed lncRNAs highlight the dynamic nature of these modifications and the need for precise frequency estimation that can inform about the functionality of a given modified site.

## DISCUSSION

Establishing a biochemical framework that allows the identification and measurable characterization of RNA epitranscriptomic modifications to complement the existing transcriptome-wide datasets, is critical for understanding RNA functionality and modes by which it is tuned in response to various environmental stimuli. The stoichiometry of modification at a given site can reflect the biological significance of that residue, as these versatile chemical tags have been shown to influence almost every aspect of RNA biology, e.g., structure, stability, metabolism, and interactions with effectors. It is currently unclear whether epitranscriptomic modifications of some target RNAs, e.g., less abundant cellular RNAs or viral transcripts, result from off-target activities of modifying cellular enzymes, or if they represent a new layer of post-transcriptional control. However, considering their extensive influence over RNA biology, they should be regarded as an additional layer of molecular code that governs specific biological effects at the cellular and organismal levels.

**Figure 4.**
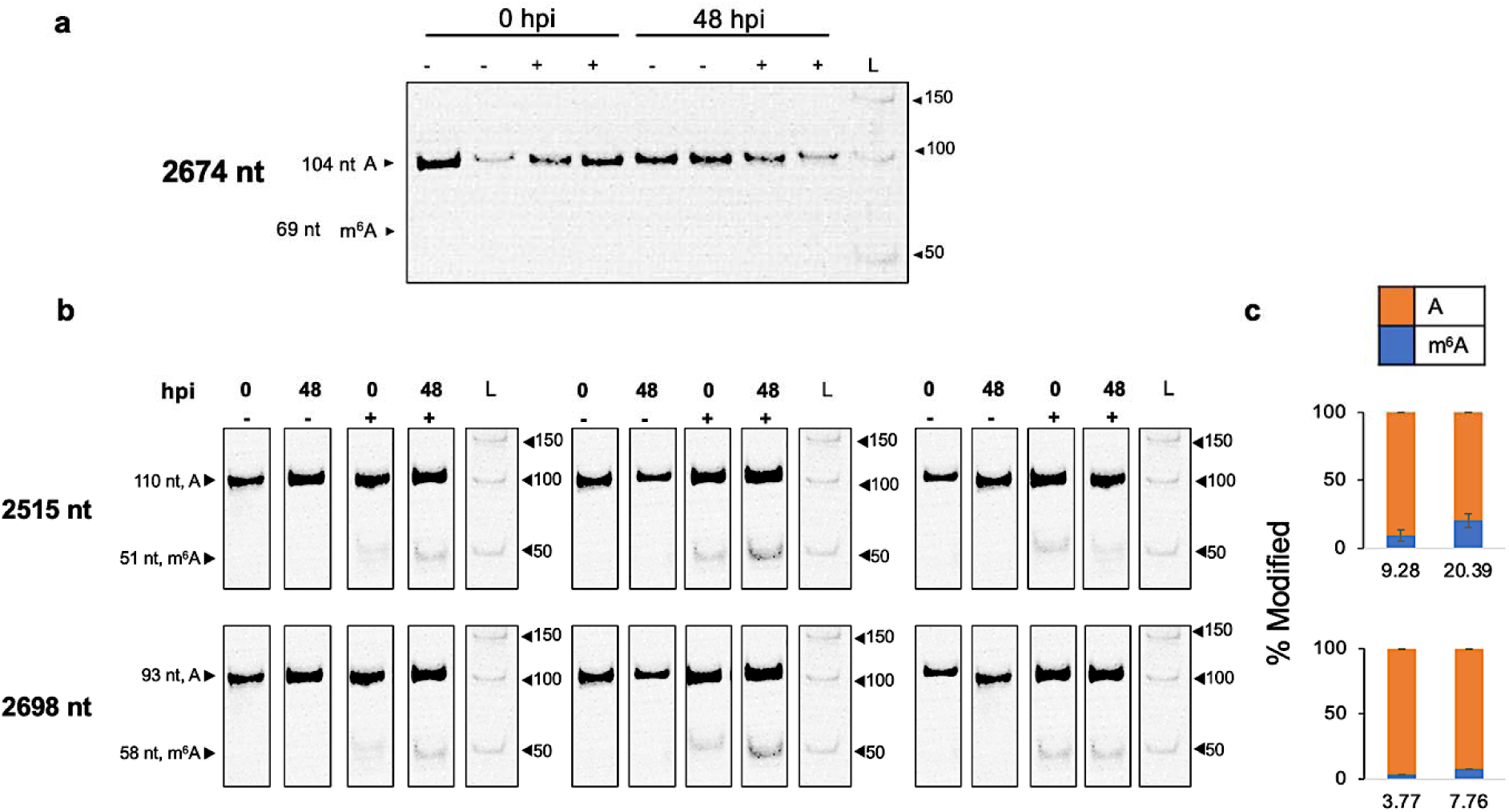
The stoichiometric analysis of m^6^A on MALAT1. **a**, Native PAGE showing one product (nts 104) corresponding to the unmodified adenine at nt position 2674 that served as a negative control. The position of expected products specific for modified and unmodified sites are indicated on all electropherograms. L stands for 50 bp DNA ladder. **b**, Native PAGE for representative three biological replicates of the SLAP analysis performed on MALAT1 for adenines at nt positions 2515 and 2698. **c**, The column graphs represent the average m^6^A modification frequency on MALAT1 RNA at nt positions 2515, and 2698. Standard deviations for frequency measurements are indicated.

Here, we present the application of a newly developed antibody-independent method, termed the SLAP, to measure the m^6^A stoichiometry on two exemplary lncRNAs, PAN RNA and MALAT1, both of which have been previously reported as m^6^A modified (Martin et al. 2021; Liu et al. 2013a). We show that the application of SLAP is not limited to a specific type of RNA in question, as we were able to obtain quantitative measurement of m^6^A levels for transcripts of viral and cellular origin. The SLAP relies on the use of site-specific oligonucleotides to “read” the modification stoichiometry regardless of its position, i.e., inside, or outside of the consensus RRACH motif. As such, we were able to determine the frequency of modification for adenines located within motifs “AAC” (nt 203 and 1041) and “CAC” (nt 18) on PAN RNA, and within motifs “GAC” (nt 2515) and “AAC” (nt 2698) on MALAT1. This is particularly critical for the analysis of m^6^A residues that were found outside the canonical consensus sequence in human (Linder et al. 2015), viral (Baquero-Perez et al. 2019), bacterial (Deng et al. 2015), and plant (Y et al. 2014; Wei et al. 2018) transcriptomes.

The SLAP method does not require large amounts of input material. Our analysis indicated that as little as 2.98 × 10^8^ copies of PAN RNA and 4 × 10^7^ copies of MALAT1 can be used to yield the accurate estimates of modification frequency on transcripts expressed in living cells. Considering the PAN RNA and MALAT1 transcripts’ average copy number per cell, this translates to approximately 600 cells. Also, using in vitro synthesized RNA standards, that we can fully control in terms of abundance and modification fraction, we were able to quantify the modification on target transcript that was present at 500 attomolar concentration (5.5 × 10^8^ copies), and the modification percentage as low as 2.5% (4.5 × 10^8^ copies of modified transcripts). For the assessment of other less abundant transcripts, one can scale up the amount of input material to obtain results of comparable accuracy.

The SLAP includes relatively small number of methodological steps that are rooted in traditional molecular biology techniques, involving total RNA extraction, phosphorylation, reverse transcription, RNA hydrolysis, ligation, PCR, and densitometric assessment of quantitative data. Further, the deconvolution of data resulting from the SLAP does not require an extensive bioinformatic background, unlike most deep-sequencing techniques. On average it takes 2 - 3 days from the isolation of the total RNA to the stoichiometric results. In comparison, other methods, e.g., m^6^A mapping by next generation sequencing methods (Linder et al. 2015; Grozhik et al. 2017; Dominissini et al. 2013; Molinie et al. 2016) are burden with many laborious steps that are required for cDNA library preparation followed by the bioinformatic analysis of obtained sequencing reads. Also, the SCARLET method, although can precisely determine m^6^A modification sites at single-nucleotide resolution, is time consuming, laborious, and it requires the use of radioactivity. Hence, the SLAP method offers an attractive alternative that has low cost and requires low time contribution or expertise.

The epitranscriptomic field is quickly opening a new chapter in RNA biology field, advancing through the discovery of novel modifications to their biological functions in many molecular processes but also human pathogenesis. It has been estimated that the defects in RNA modifications account for over 100 human disorders that include childhood-onset multiorgan failures, cancers, metabolic, and neurologic diseases (Suzuki 2020; Jonkhout et al. 2017). These conditions are now referred to as “RNA modopathies”, and the extent to which their severity is defined by the disruption of epitranscriptomic processes is under careful examination. The next horizon for this quickly progressing field is to establish a molecular level view of how these chemical tags define their influence over single transcript, whole transcriptome, cellular processes, and phenotypic consequences. We now realize that RNA sequences and structures with their modifications comprise the complete information content of the RNA. They are needed to usher in an era of molecular and clinical studies that are based on a solid foundation of sequences and structures.

## MATERIALS & METHODS

### Oligonucleotide Design

The analysis of each modified adenine requires the design of four oligonucleotides: 1) an oligonucleotide primer with a minimum length of 20 nts and matching the region located at least 50 nts downstream of the target m^6^A. This reverse primer (Figure 5a) binds the modified RNA template and serves as a starting point for synthesis of a new copy DNA (cDNA) during reverse transcription (RT) reaction. Since RT reaction takes place in the presence of 4SedTTP, it can yield two types of products: a truncated “RT-stop” product marking the position of m^6^A, and a full-length product corresponding to the unmodified RNA fraction (Figure 5b). 2) the forward primer (Figure 5a) with a minimum length of 20 nts, that holds complementarity to the region located at least 25 nts upstream of the target m^6^A, which allows the simultaneous amplification of both types of products. 3) the 5’ phosphorylated adapter oligonucleotide containing CCATTG insert at the 5’ end (Figure 5c), while its 3’ end contains sequence that is reverse complement to the forward primer binding site. 4) the splint oligonucleotide (Figure 5c), which 5’ end is reverse complement to the adapter sequence necessary for duplex formation (including the adapter’s CCATTG insert, and forward primer binding site), followed by the sequence that is reverse compliment to the 10 nts downstream of the target m^6^A site (not including the modified residue), and the 3’ end, that includes the 3’ C3 spacer phosphoramidite (SpC3). The SpC3 modification introduces a long hydrophilic spacer arm for the ligation of the phosphorylated adapter. As a result of ligation reaction, both RT-stop and full-length products will include common reverse and forward primer binding sites that support their simultaneous PCR amplification. Altogether, the total length of ligated adapter, splint and RT-stop product should not approximate the size of full-length product to allow the size-specific discrimination; thus, we recommend at least 25 nt difference between both. For target adenines that are located near the 5’ end of modified RNA (within 50 nts, as for m^6^A at position 18 in PAN RNA), the design of extended splint oligonucleotides is required to adequately differentiate between RT-stop and full-sized products. This can be accomplished by the addition of non-template specific sequence to the adapter oligonucleotide that will follow the CCATTG insert and proceed the 3’ end sequence that is reverse complement to the forward primer.

**Figure 5.**
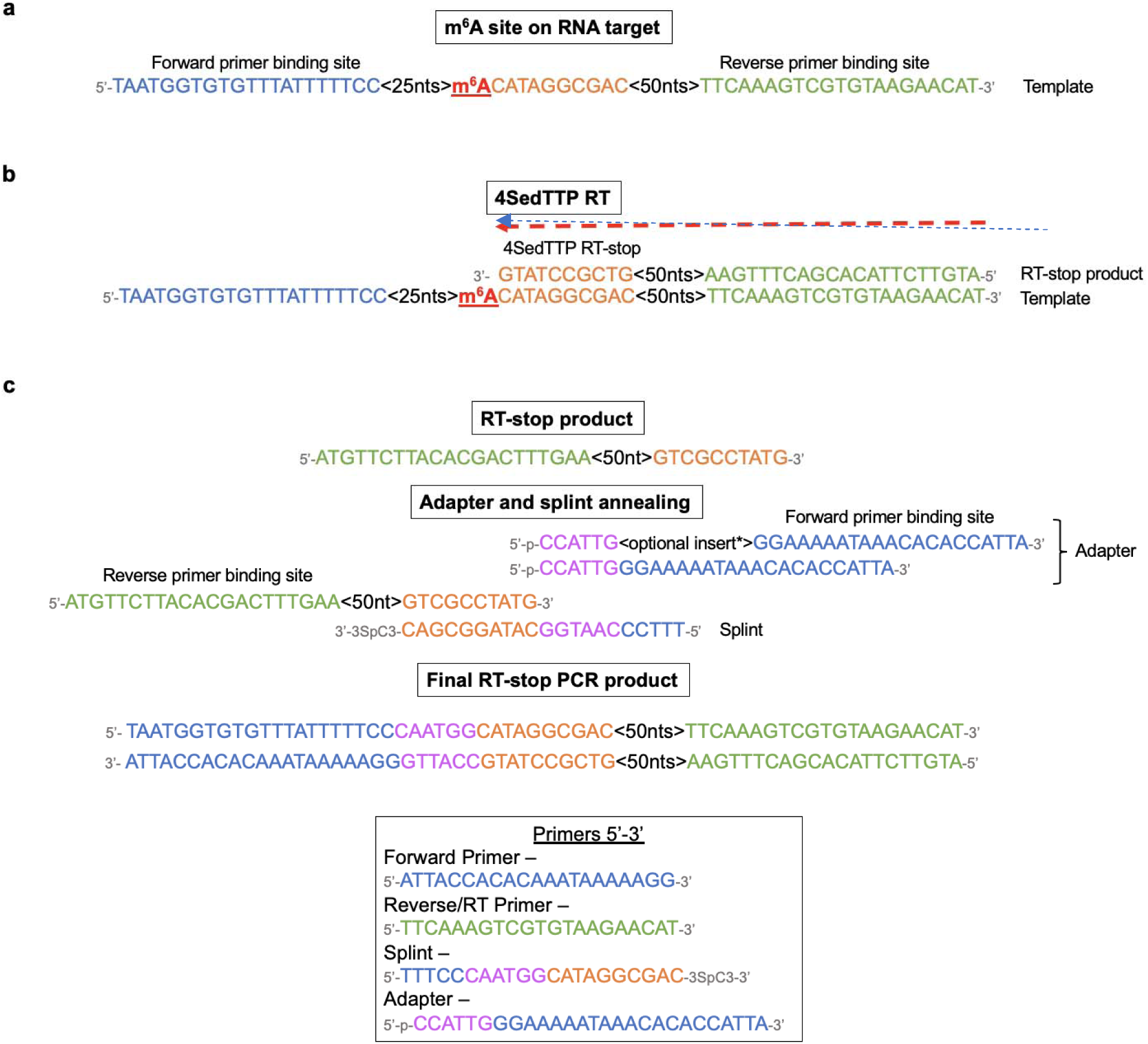
The guidelines for the design of SLAP primers. **a**, The representative sequence of modified target RNA with indicated m^6^A (in red). 25 and 50 nts sections on the target RNA refer to the number of nucleotides required for spacing between forward primer binding site (in blue) and reverse primer binding site (in green). Orange sequence corresponds to 10 nts downstream of the m^6^A site, which acts as a sticky end for splint bridge annealing. **b**, The m^6^A modified RNA is directed to 4SedTTP reverse transcription during which the presence of 4SedTTP induces RT-stop at +1 position from m^6^A. **c**, The RT-stop product is ligated to splint-adapter oligonucleotide duplex, with adapter including forward primer binding site and a splint including the 3SpC3 modification at its 3’ end. For m^6^A that are near 5’ end, an extended adapter oligonucleotide should be used to allow size-specific differentiation between RT-stop and full-size products corresponding to modified and unmodified RNA, respectively.

### Cell lines and culture conditions

The KSHV positive body cavity-based lymphoma cell line (BCBL-1, a generous gift from Dr. Denise Whitby, NCI Frederick) was seeded at 2 × 10^5^ cells/ml and grown in RPMI-1640 medium (ThermoFisher 11875085) supplemented with 10% fetal bovine serum (FBS), 1% Penicillin/Streptomycin and 1% L-Glutamine at 37 °C in 5% CO_2_. The induction of KSHV lytic infection was performed by treating 2 × 10^7^ BCBL-1 cells with sodium butyrate (NaB, EMD Millipore 654833) to a final concentration of 0.3 mM. Cells were collected before (0 h post induction, h pi) and after induction (48 h pi).

### RNA extraction

Total RNA was isolated from 2×10^7^ BCBL-1 cells using TRIzol™ (ThermoFisher 15596026), followed by DNase I treatment (ThermoFisher AM1907), and RNA purification using RNA Clean & Concentrator-5. RNA was eluted in of RNase/DNase-free water and stored at -80°C.

### 5’ Phosphorylation of total RNA

The 5’ end phosphorylation was performed by treating 10 μg of total RNA extract with 0.5 μl of RNase inhibitor (20U final, NEB M0307L), 1 μl of 10x T4 PNK reaction buffer, 1 μl 0.1 mM ATP, and 1 μl of T4 Polynucleotide kinase (10U final) in 10 μl total volume for 30 min at 37 °C. RNA was ethanol precipitation by adding 160 μl of 50 mM KOAc, pH 7, 200 mM KCl, 3 μl of 5 μg/μl Glycogen, and 550 μl 100% ethanol and stored at -20°C. The samples were centrifuged at 12,000 x g for 30 minutes, the pellet was washed with 500 μl of 75% ethanol and dissolved in 13.5 μl of RNase/DNase-free water.

### 4SedTTP reverse transcription

The reverse transcription reaction included 13.5 μl of the phosphorylation reaction that was initially combined with 1 μl 10x annealing buffer (250 mM Tris-HCl, pH 7.4, 480 mM KCl), and 1 μl 0.5 pmol RT oligonucleotide primer (Table 1) in the total volume of 15.5 μl. The reaction was incubated for 2 min at 95°C. For 4SedTTP reactions, 2 µl of 10X 4SedTTP reaction buffer (500 mM Tris-HCl pH 8.0, 500 mM KCl, 50 mM MgCl_2_, 100 mM DTT), 1 µl of 800 µM dATP, dCTP, dGTP, 4SedTTP (final concentration of 80 µM for each), 0.5 µl RNase Inhibitor (80U final, NEB M0307L) and 1 µl AMV RT (12U final, ThermoFisher 18080044) were added to each reaction for a final volume of 20 µl. The reactions were incubated for 1 h at 42°C and 5 min at 85°C to inactivate the reverse transcriptase. To remove RNA template, 1 µl RNase H (5U final, NEB M0297L) was added directly to each reaction and incubation for 20 min at 37°C. RNase H was inactivated by incubating for 5 min at 85°C. For dTTP reactions, the above protocol was performed except for the replacement of 4SedTTP with 800 µM dTTP (80 µM final concentration) during RT reactions

**Table 1.**
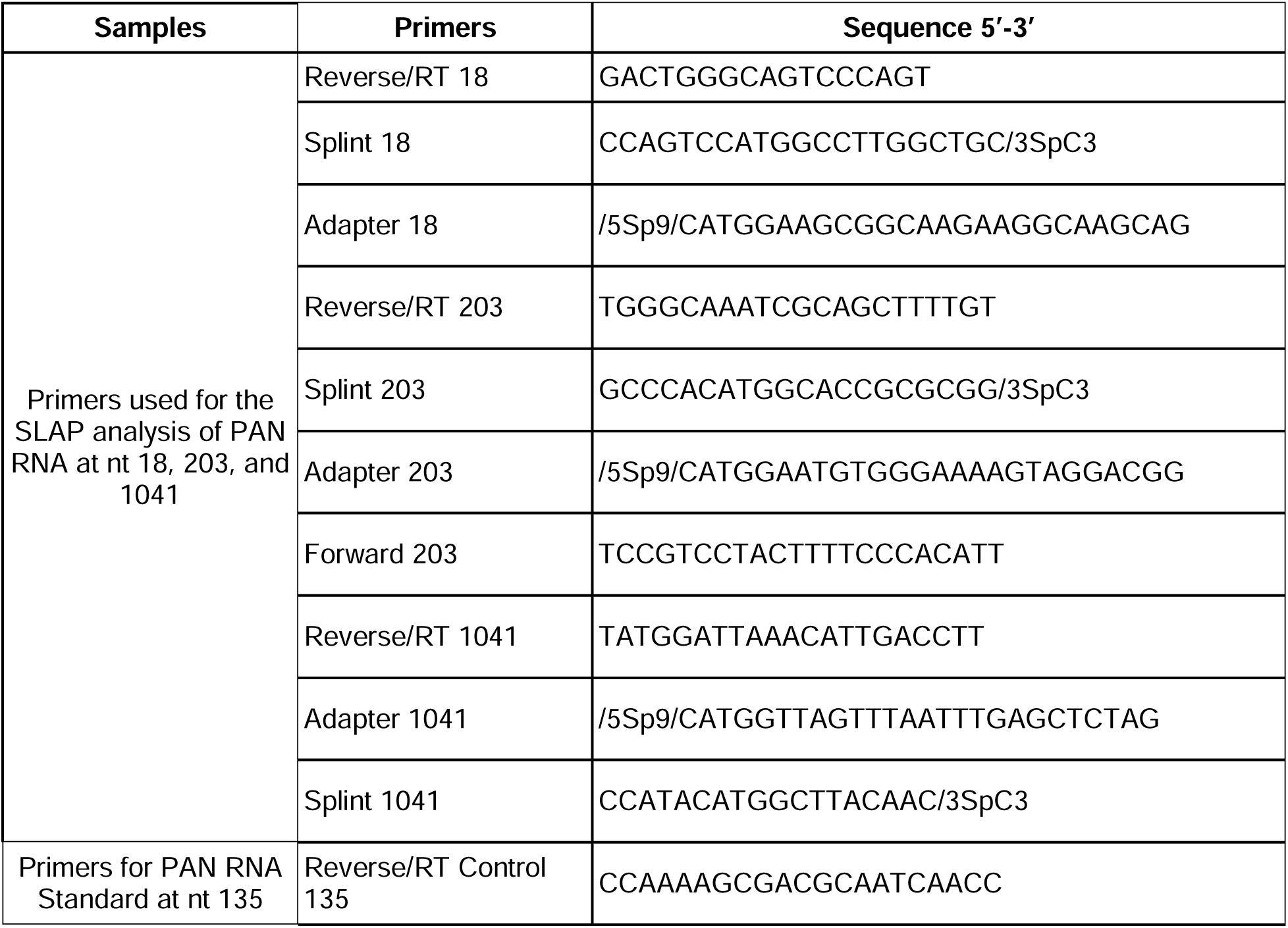
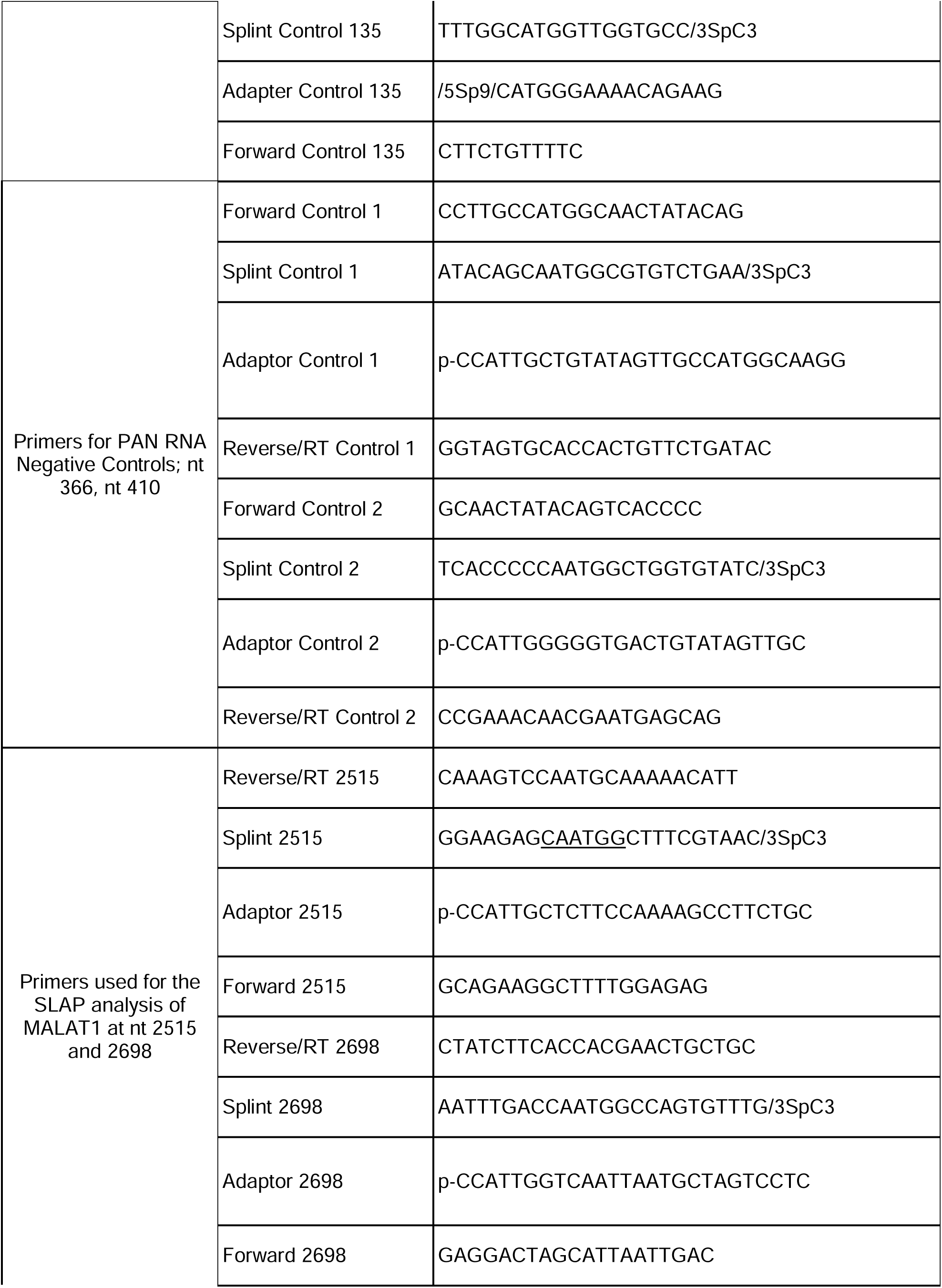
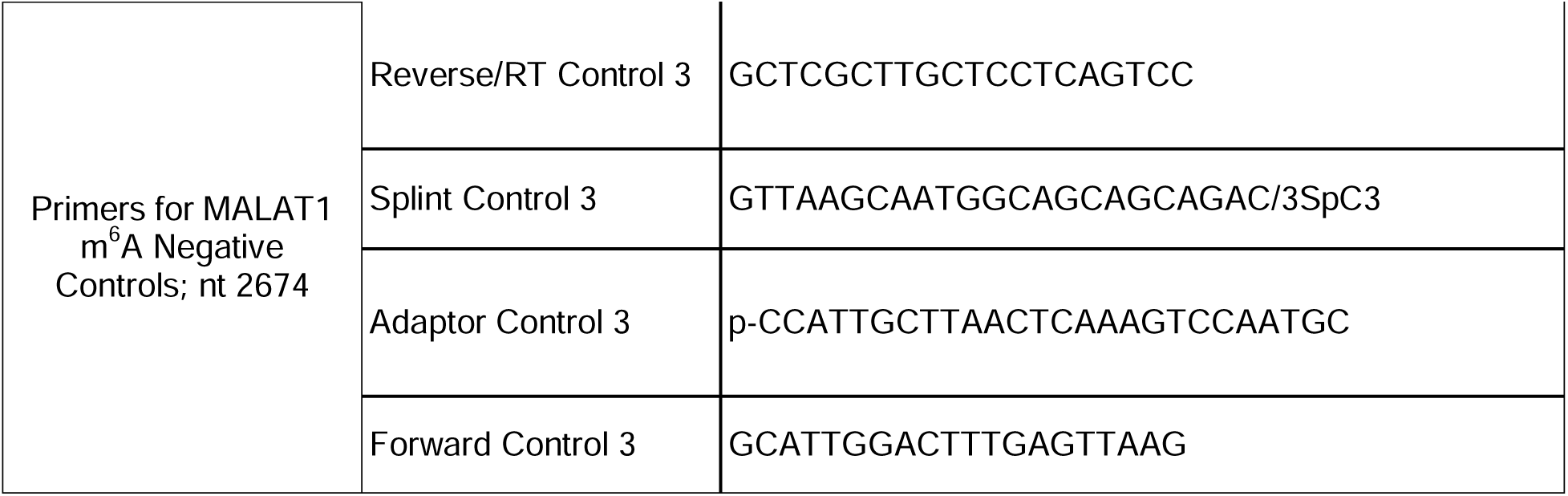
The SLAP oligonucleotides.

### Annealing and ligation

The adapters and splint oligonucleotides were combined at 1:1 ratio, at a concentration of 1.5 μM each. 1.5 pmol (1 μl) of the adapter/splint oligonucleotides mixture was added to 20 μl RT reaction followed by incubation for 3 min at 75°C to facilitate the annealing of splint oligonucleotide to RT-stop products and the adapter to the splint oligonucleotide. Next, to ligate the adapter to the RT-stop product, 0.4 μl of 10 U/μl T4 DNA ligase (4U final, NEB M0202L), 1X T4 DNA ligase reaction buffer, and 2 μl 100% DMSO were added to the total volume of 25 μl and incubated overnight at 16 °C. The DNA ligase was inactivated by incubating the reaction for 10 min at 65°C.

### PCR Amplification of RT-stop and full-length products

To amplify both RT-stop and full-length products, 2 μl of the ligation reaction was combined with 10 μl HiFi Buffer (-Mg^2+^), 2 μl 10 mM dNTPs, 4 μl 50 mM MgCl_2_, 2 μl 10 μM forward primer, 2 μl 10 μM RT/reverse primer, 71.6 μl water, 0.4 μl Platinum Taq High Fidelity DNA Polymerase (2U final, ThermoFisher 11304011) in 25 μl total. The reaction was initially denatured for 30s at 94°C, followed by 35 cycles of denaturation for 10s at 94°C, annealing for 20s at 67°C, and extension for 20s at 72°C. The final extension was set for 5 min at 72°C.

### Native polyacrylamide gel electrophoresis and densitometric analysis

The PCR products were separated on 10% native polyacrylamide gel (PAGE) that was set by combining 9.67 ml RNase/DNase-free water, 1.5 ml 10x Tris-Borate-EDTA buffer (10x TBE, pH 8.3), 3.75 ml 40% Acrylamide/Bisacrylamide (19:1), 75 μl 10% ammonium persulfate, and 7.5 μl N,N,N’,N’-tetramethylethylenediamine (TEMED). After combining samples with a gel loading buffer, which consisted of 30% (v/v) glycerol, and 0.25% (w/v) bromophenol blue, the samples were loaded on the gel. The PAGE was run for 2 h at 120V in 1X TBE buffer. The gel was stained by incubating with 1x ethidium bromide solution in 1x TBE overnight at 4 °C (Zhang et al. 2019). Gels were visualized using the Benchtop 3UV Transilluminator and GelDoc-IT Imager. Densitometric analysis was performed using ImageJ v1.52a (Abràmoff et al. 2006) on inverted TIFF files. Rectangular boxes were drawn around each band, and the measure function was used to determine their pixel density. Background was determined by densitometry measurements of three randomly chosen areas located near the area where the RT-stop products would be expected in (-)4SedTTP reactions. This measurement was subtracted from the experimental pixel density measurements. The stoichiometric measurements of m^6^A were visualized as bar graphs using Excel spreadsheet. Measured values corresponding to RT-stop and full-length PCR products were used to calculate percent yield using the following formula:

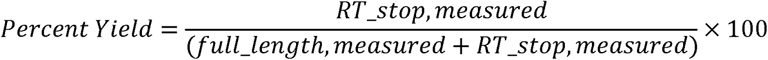

## Acknowledgement

Research reported in this publication and S.E.M., H.G., and J.S.S. were supported by the start-up funds from the Department of Biological Sciences, College of Science and Mathematics, and Office of the Vice President for Research, Auburn University, and by the National Institute of Allergy and Infectious Disease (NIAID) under award number R21AI159361. The content is solely the responsibility of the authors and does not necessarily represent the official views of the National Institutes of Health.

## REFERENCES

Abràmoff MD, Magalhães PJ, Ram SJ. 2006. Image Processing with ImageJ. In Optical Imaging Techniques in Cell Biology, pp. 249–258, CRC Press.

Arun G, Aggarwal D, Spector DL. 2020. MALAT1 Long Non-Coding RNA: Functional Implications. Non-Coding RNA 6. /pmc/articles/PMC7344863/ (Accessed September 14, 2021).

Baquero-Perez B, Antanaviciute A, Yonchev ID, Carr IM, Wilson SA, Whitehouse A. 2019. The tudor SND1 protein is an m6a RNA reader essential for replication of kaposi’s sarcoma-associated herpesvirus. eLife 8.

Baquero-Perez B, Geers D, Díez J. 2021. From A to m6A: The Emerging Viral Epitranscriptome. Viruses 13: 1049. https://www.mdpi.com/1999-4915/13/6/1049 (Accessed June 24, 2021).

Bartosovic M, Molares HC, Gregorova P, Hrossova D, Kudla G, Vanacova S. 2017. N6-methyladenosine demethylase FTO targets pre-mRNAs and regulates alternative splicing and 3’-end processing. Nucleic acids research 45: 11356–11370. http://www.ncbi.nlm.nih.gov/pubmed/28977517 (Accessed October 30, 2019).

Bokar JA. 2005. The biosynthesis and functional roles of methylated nucleosides in eukaryotic mRNA. 141–177. https://link.springer.com/chapter/10.1007/b106365 (Accessed July 13, 2021).

Bokar JA, Shambaugh ME, Polayes D, Matera AG, Rottman FM. 1997. Purification and cDNA cloning of the AdoMet-binding subunit of the human mRNA (N6-adenosine)-methyltransferase. RNA 3: 1233–1247. http://www.rnajournal.org/cgi/content/abstract/3/11/1233#otherarticles (Accessed August 30, 2020).

Coker H, Wei G, Brockdorff N. 2019. m6A modification of non-coding RNA and the control of mammalian gene expression. Biochimica et Biophysica Acta - Gene Regulatory Mechanisms 1862: 310–318.

Deng X, Chen K, Luo G-Z, Weng X, Ji Q, Zhou T, He C. 2015. Widespread occurrence of N6-methyladenosine in bacterial mRNA. Nucleic Acids Research 43: 6557. /pmc/articles/PMC4513869/ (Accessed September 15, 2021).

Dominissini D, Moshitch-Moshkovitz S, Salmon-Divon M, Amariglio N, Rechavi G. 2013. Transcriptome-wide mapping of N6-methyladenosine by m6A-seq based on immunocapturing and massively parallel sequencing. Nature Protocols 2013 8:1 8: 176–189. https://www.nature.com/articles/nprot.2012.148 (Accessed September 14, 2021).

Dominissini D, Moshitch-Moshkovitz S, Schwartz S, Salmon-Divon M, Ungar L, Osenberg S, Cesarkas K, Jacob-Hirsch J, Amariglio N, Kupiec M, et al. 2012. Topology of the human and mouse m6A RNA methylomes revealed by m6A-seq. Nature 485: 201–206. https://www.nature.com/articles/nature11112 (Accessed May 12, 2021).

Grozhik A v., Linder B, Olarerin-George AO, Jaffrey SR. 2017. Mapping m6A at individual-nucleotide resolution using crosslinking and immunoprecipitation (miCLIP). Methods in molecular biology (Clifton, NJ) 1562: 55. /pmc/articles/PMC5562447/ (Accessed July 13, 2021).

He RZ, Jiang J, Luo DX. 2020. The functions of N6-methyladenosine modification in lncRNAs. Genes & Diseases 7: 598–605.

Hong T, Yuan Y, Chen Z, Xi K, Wang T, Xie Y, He Z, Su H, Zhou Y, Tan ZJ, et al. 2018. Precise Antibody-Independent m6A Identification via 4SedTTP-Involved and FTO-Assisted Strategy at Single-Nucleotide Resolution. Journal of the American Chemical Society 140: 5886–5889. https://pubs.acs.org/sharingguidelines (Accessed October 30, 2019).

Jiang X, Liu B, Nie Z, Duan L, Xiong Q, Jin Z, Yang C, Chen Y. 2021. The role of m6A modification in the biological functions and diseases. Signal Transduction and Targeted Therapy 6: 1–16. https://doi.org/10.1038/s41392-020-00450-x (Accessed May 2, 2021).

Jonkhout N, Tran J, Smith MA, Schonrock N, Mattick JS, Novoa EM. 2017. The RNA modification landscape in human disease. RNA 23: 1754. /pmc/articles/PMC5688997/ (Accessed September 14, 2021).

Kane SE, Beemon K. 1985. Precise localization of m6A in Rous sarcoma virus RNA reveals clustering of methylation sites: implications for RNA processing. Molecular and Cellular Biology 5: 2298–2306. https://pubmed.ncbi.nlm.nih.gov/3016525/ (Accessed September 16, 2020).

KD M, Y S, P Z, O E, CE M, SR J. 2012. Comprehensive analysis of mRNA methylation reveals enrichment in 3’ UTRs and near stop codons. Cell 149: 1635–1646. https://pubmed.ncbi.nlm.nih.gov/22608085/ (Accessed September 14, 2021).

Linder B, Grozhik A v., Olarerin-George AO, Meydan C, Mason CE, Jaffrey SR. 2015. Single-nucleotide-resolution mapping of m6A and m6Am throughout the transcriptome. Nature Methods 12: 767–772. https://www.nature.com/articles/nmeth.3453 (Accessed May 12, 2021).

Liu J, Yue Y, Han D, Wang X, Fu Y, Zhang L, Jia G, Yu M, Lu Z, Deng X, et al. 2014. A METTL3-METTL14 complex mediates mammalian nuclear RNA N6-adenosine methylation. Nature chemical biology 10: 93–95. https://www.nature.com/articles/nchembio.1432 (Accessed August 20, 2020).

Liu N, Parisien M, Dai Q, Zheng G, He C, Pan T. 2013a. Probing N6-methyladenosine RNA modification status at single nucleotide resolution in mRNA and long noncoding RNA. RNA 19: 1848. /pmc/articles/PMC3884656/ (Accessed September 4, 2021).

Liu N, Parisien M, Dai Q, Zheng G, He C, Pan T. 2013b. Probing N6-methyladenosine RNA modification status at single nucleotide resolution in mRNA and long noncoding RNA. RNA 19: 1848–1856. /pmc/articles/PMC3884656/?report=abstract (Accessed September 7, 2020).

Lu Z, Guo JK, Wei Y, Dou DR, Zarnegar B, Ma Q, Li R, Zhao Y, Liu F, Choudhry H, et al. 2020. Structural modularity of the XIST ribonucleoprotein complex. Nature Communications 2020 11:1 11: 1–14. https://www.nature.com/articles/s41467-020-20040-3 (Accessed July 20, 2021).

Mao Y, Dong L, Liu X-M, Guo J, Ma H, Shen B, Qian S-B. 2019. m 6 A in mRNA coding regions promotes translation via the RNA helicase-containing YTHDC2. Nature Communications 2019 10:1 10: 1–11. https://www.nature.com/articles/s41467-019-13317-9 (Accessed July 20, 2021).

Martin SE, Gan H, Toomer G, Sridhar N, Sztuba-Solinska J. 2021. The m6A landscape of polyadenylated nuclear (PAN) RNA and its related methylome in the context of KSHV replication. RNA rna.078777.121. http://rnajournal.cshlp.org/content/early/2021/06/29/rna.078777.121 (Accessed July 13, 2021).

Mateusz Mendel A, Chen K-M, Homolka D, Gos P, Raman Pandey R, McCarthy AA, Pillai RS, Mendel M. 2018. Methylation of Structured RNA by the m 6 A Writer METTL16 Is Essential for Mouse Embryonic Development Article Methylation of Structured RNA by the m 6 A Writer METTL16 Is Essential for Mouse Embryonic Development Internal modification of RNAs with N 6. Molecular Cell 71: 986–1000.

McIntyre W, Netzband R, Bonenfant G, Biegel JM, Miller C, Fuchs G, Henderson E, Arra M, Canki M, Fabris D, et al. 2018. Positive-sense RNA viruses reveal the complexity and dynamics of the cellular and viral epitranscriptomes during infection. Nucleic Acids Research 46: 5776–5791. https://pubmed.ncbi.nlm.nih.gov/29373715/ (Accessed September 16, 2020).

Mendel M, Delaney K, Pandey RR, Chen K-M, Wenda JM, Vågbø CB, Steiner FA, Homolka D, Pillai RS. 2021. Splice site m6A methylation prevents binding of U2AF35 to inhibit RNA splicing. Cell 184: 3125-3142.e25.

Meyer KD, Patil DP, Zhou J, Zinoviev A, Skabkin MA, Elemento O, Pestova T v., Qian SB, Jaffrey SR. 2015. 51 UTR m6A Promotes Cap-Independent Translation. Cell 163: 999–1010.

Meyer KD, Saletore Y, Zumbo P, Elemento O, Mason CE, Jaffrey SR. 2012. Comprehensive analysis of mRNA methylation reveals enrichment in 31 UTRs and near stop codons. Cell 149: 1635–1646. https://pubmed.ncbi.nlm.nih.gov/22608085/ (Accessed June 17, 2021).

Molinie B, Wang J, Lim KS, Hillebrand R, Lu Z, van Wittenberghe N, Howard BD, Daneshvar K, Mullen AC, Dedon P, et al. 2016. m6A-LAIC-seq reveals the census and complexity of the m6A epitranscriptome. Nature Methods 2016 13:8 13: 692–698. https://www.nature.com/articles/nmeth.3898 (Accessed September 4, 2021).

Rossetto CC, Pari GS. 2014. PAN’s labyrinth: Molecular biology of kaposi’s sarcoma-associated herpesvirus (KSHV) PAN RNA, a multifunctional long noncoding RNA. Viruses 6: 4212–4226.

Schwartz S, Mumbach MR, Jovanovic M, Wang T, Maciag K, Bushkin GG, Mertins P, Ter-Ovanesyan D, Habib N, Cacchiarelli D, et al. 2014. Perturbation of m6A writers reveals two distinct classes of mRNA methylation at internal and 5’ sites. Cell Reports 8: 284–296. /pmc/articles/PMC4142486/?report=abstract (Accessed August 30, 2020).

Śledź P, Jinek M. 2016. Structural insights into the molecular mechanism of the m6A writer complex. eLife 5. /pmc/articles/PMC5023411/ (Accessed July 20, 2021).

Suzuki T. 2020. RNA Modifications in Health and Disease. The FASEB Journal 34: 1–1. https://onlinelibrary.wiley.com/doi/full/10.1096/fasebj.2020.34.s1.00132 (Accessed September 14, 2021).

Tan B, Gao S-J. 2018. The RNA Epitranscriptome of DNA Viruses. Journal of Virology 92. https://pubmed.ncbi.nlm.nih.gov/30185592/ (Accessed December 14, 2020).

Tripathi V, Ellis JD, Shen Z, Song DY, Pan Q, Watt AT, Freier SM, Bennett CF, Sharma A, Bubulya PA, et al. 2010. The Nuclear-Retained Noncoding RNA MALAT1 Regulates Alternative Splicing by Modulating SR Splicing Factor Phosphorylation. Molecular cell 39: 925. /pmc/articles/PMC4158944/ (Accessed September 14, 2021).

Wang X, Feng J, Xue Y, Guan Z, Zhang D, Liu Z, Gong Z, Wang Q, Huang J, Tang C, et al. 2016. Structural basis of N6-adenosine methylation by the METTL3-METTL14 complex. Nature 534: 575–578.

Wang X, Lu Z, Gomez A, Hon GC, Yue Y, Han D, Fu Y, Parisien M, Dai Q, Jia G, et al. 2014. N 6-methyladenosine-dependent regulation of messenger RNA stability. Nature 505: 117–120. https://www.nature.com/articles/nature12730 (Accessed June 17, 2021).

Wang X, Zhao BS, Roundtree IA, Lu Z, Han D, Ma H, Weng X, Chen K, Shi H, He C. 2015. N6-methyladenosine modulates messenger RNA translation efficiency. Cell 161: 1388–1399.

Wei L-H, Song P, Wang Y, Lu Z, Tang Q, Yu Q, Xiao Y, Zhang X, Duan H-C, Jia G. 2018. The m6A Reader ECT2 Controls Trichome Morphology by Affecting mRNA Stability in Arabidopsis. The Plant Cell 30: 968–985. https://academic.oup.com/plcell/article/30/5/968/6099257 (Accessed September 15, 2021).

Wein S, Andrews B, Sachsenberg T, Santos-Rosa H, Kohlbacher O, Kouzarides T, Garcia BA, Weisser H. 2020. A computational platform for high-throughput analysis of RNA sequences and modifications by mass spectrometry. Nature Communications 11: 1–12. https://doi.org/10.1038/s41467-020-14665-7 (Accessed May 10, 2021).

Withers JB, Li ES, Vallery TK, Yario TA, Steitz JA. 2018. Two herpesviral noncoding PAN RNAs are functionally homologous but do not associate with common chromatin loci. PLoS Pathogens 14. /pmc/articles/PMC6233925/?report=abstract (Accessed December 9, 2020).

Wu F, Cheng W, Zhao F, Tang M, Diao Y, Xu R. 2019. Association of N6-methyladenosine with viruses and related diseases. Virology Journal 16. /pmc/articles/PMC6849232/ (Accessed July 13, 2021).

Y L, X W, C L, S H, J Y, S S. 2014. Transcriptome-wide N<sup>^6^</sup>-methyladenosine profiling of rice callus and leaf reveals the presence of tissue-specific competitors involved in selective mRNA modification. RNA biology 11: 1180–1188. https://pubmed.ncbi.nlm.nih.gov/25483034/ (Accessed September 15, 2021).

Yang F, Yi F, Han X, Du Q, Liang Z. 2013. MALAT-1 interacts with hnRNP C in cell cycle regulation. FEBS Letters 587: 3175–3181. https://pubmed.ncbi.nlm.nih.gov/23973260/ (Accessed March 28, 2021).

Yang Y, Hsu PJ, Chen YS, Yang YG. 2018. Dynamic transcriptomic m6A decoration: Writers, erasers, readers and functions in RNA metabolism. Cell Research 28: 616–624. https://pubmed.ncbi.nlm.nih.gov/29789545/ (Accessed December 9, 2020).

Zhang W, Eckwahl MJ, Zhou KI, Pan T. 2019. Sensitive and quantitative probing of pseudouridine modification in mRNA and long noncoding RNA. RNA 25: 1218–1225.

Zhou KI, Shi H, He C, Parisien M. 2019a. Regulation of Co-transcriptional Pre-mRNA Splicing by m 6 A through the Low-Complexity Protein hnRNPG. Molecular Cell 76: 70–81. https://doi.org/10.1016/j.molcel.2019.07.005 (Accessed May 12, 2021).

Zhou KI, Shi H, Lyu R, Wylder AC, Matuszek Ż, Pan JN, He C, Parisien M, Pan T. 2019b. Regulation of Cotranscriptional Pre-mRNA Splicing by m6A through the Low-Complexity Protein hnRNPG. Molecular Cell 76: 70-81.e9. https://pubmed.ncbi.nlm.nih.gov/31445886/ (Accessed September 16, 2020).

